## Supplementary Table 1 for "Locus-specific enrichment analysis of 5-hydroxymethylcytosine reveals novel genes associated with breast carcinogenesis"

### ST. 1a. Genomic Feature of 5-mC in breast cancer and normal tissues

| Genomic Features | | Tumour (IDC) (n>=10) (%) | Paired Normal (n>=10) (%) | DCIS (n=5) (%) | Apparent normal (n=5) (%) |
|---|---|---|---|---|---|
| **Promoters** | Promoter(>1kb) | 8.24 | 6.09 | 7.41 | 6.34 |
| | Promoter(1-2kb) | 4.17 | 3.44 | 3.76 | 3.3 |
| | Promoter(2-3kb) | 3.68 | 3.15 | 3.31 | 2.88 |
| **UTRs** | 5'UTR | 1.69 | 1.32 | 1.54 | 1.4 |
| | 3'UTR | 4.35 | 3.3 | 3.89 | 3.32 |
| **Gene body** | 1st exon | 0.51 | 0.4 | 0.46 | 0.42 |
| | Other exon | 9.41 | 7.98 | 8.52 | 7.72 |
| | 1st intron | 9.72 | 9.7 | 9.44 | 9.57 |
| | Other intron | 24.26 | 24.64 | 24.42 | 24.43 |
| **Distal intergenic** | Downstream (>300kb) | 0.16 | 0.12 | 0.12 | 0.12 |
| | Distal intergenic | 33.81 | 39.86 | 37.13 | 40.5 |

### ST. 1b. Genomic Feature of 5-hmC in breast cancer and normal tissues

| Genomic features | | Tumour (IDC) (n>=10) (%) | Paired Normal (n>=10) (%) | DCIS (n=5) (%) | Apparent normal (n=5) (%) |
|---|---|---|---|---|---|
| **Promoters** | Promoter(>1kb) | 24.89 | 30.89 | 48.74 | 6.25 |
| | Promoter(1-2kb) | 2.46 | 2.3 | 1.88 | |
| | Promoter(2-3kb) | 2.6 | 2.69 | 2.06 | 6.25 |
| **UTRs** | 5'UTR | 1.46 | 1.57 | 0.97 | |
| | 3'UTR | 3.64 | 3.99 | 2.64 | |
| **Gene body** | 1st exon | 0.9 | 0.88 | 1.12 | 12.5 |
| | Other exon | 5.8 | 6.52 | 3.44 | 25 |
| | 1st intron | 7.73 | 6.63 | 4.61 | 6.25 |
| | Other intron | 14.37 | 12.66 | 7.02 | 18.75 |
| **Distal intergenic** | Downstream (>300kb) | 0.08 | 0.08 | 0.13 | |
| | Distal intergenic | 35.98 | 31.8 | 27.4 | 25 |
